# Spatially resolved mapping of tau amplification rates via differentiable simulation of prion-like propagation

**DOI:** 10.64898/2026.06.02.729568

**Authors:** Yohei Kondo, Honda Naoki, the Alzheimer’s Disease Neuroimaging Initiative

**Affiliations:** Center for One Medicine Innovative Translational Research (COMIT), Nagoya University, Nagoya, Aichi, Japan; Department of Integrative Cellular Informatics, Center for Neurological Diseases and Cancer, Nagoya University Graduate School of Medicine, Nagoya, Aichi, Japan; Graduate School of Integrated Sciences for Life, Hiroshima University, Higashihiroshima, Hiroshima, Japan

## Abstract

Neurodegenerative diseases exhibit characteristic yet heterogeneous patterns of pathological spread, whose underlying determinants remain unclear. A central challenge is that inferring spatially heterogeneous propagation kinetics from neuroimaging data constitutes a high-dimensional inverse problem that has remained intractable at the whole-brain scale. Here, we present a differentiable reaction-diffusion framework that enables inference of spatially resolved tau amplification rates from tau PET data. By integrating MRI-informed forward simulation with error backpropagation, our approach reconstructs subject-specific voxel-wise maps of tau amplification rates across the human brain. Analysis of the inferred maps showed that the amplification rates have a positive spatial association with amyloid PET, suggesting that they capture aspects of regional vulnerability to amyloid-driven tau accumulation. Furthermore, integration with transcriptomic data identified gene expression programs associated with regional variation in amplification. These findings provide a data-driven framework linking molecular architecture to large-scale propagation dynamics in neurodegeneration.

## Introduction

Neurodegenerative diseases show characteristic brain-wide patterns of pathological spread, yet their progression exhibits substantial regional and inter-individual variability [1]. In Alzheimer’s disease, neuroimaging of tau pathology indicates that the overall expansion of lesions follows anatomical connectivity, although structural constraints alone account for only part of the regional distribution of pathology [2, 3]. Furthermore, the timing and rate of regional involvement differ markedly between subjects [4]. The dynamical and molecular determinants underlying this heterogeneity remain unclear.

Mathematical modeling has been used to simulate the brain-wide spread of pathology based on a prion-like mechanism. In this framework, pathological proteins template the conformational conversion of their native counterparts and propagate between cells. Connectome-based models, built on brain parcellations, simulate the spread of abnormal proteins along inter-regional connections [5, 6]. Such connectome-based models have been further parameterized using molecular and neuroimaging data, improving their predictive performance and providing insights into the biological determinants of prion-like propagation [7, 8, 9]. In parallel, continuous spatial reaction-diffusion models have been introduced to capture finer spatial detail [10, 11, 12]. Despite their strong potential, the application of these spatially resolved models remains limited, and their ability to faithfully capture the underlying biology has yet to be fully realized. A fundamental challenge is the lack of quantitative methods to determine the spatial heterogeneity of the physical and chemical parameters governing protein propagation. As a result, model parameters are often specified in a simplistic and largely heuristic manner, limiting their ability to quantitatively reproduce disease progression. This limitation has hindered the broader application of spatial models in data-driven analyses.

Inferring spatially resolved propagation kinetics directly from neuroimaging data poses a formidable inverse problem [13]. Allowing diffusion and reaction rates to vary at the voxel level introduces more than one million unknown parameters in a whole-brain model, making brute-force optimization intractable. Error backpropagation provides an effective strategy to circumvent this curse of dimensionality. However, standard numerical solvers for partial differential equation (PDE) models of spatially continuous dynamics are not designed for gradient-based learning, preventing efficient backpropagation of errors through the dynamical system. These technical barriers have thus far precluded large-scale, data-driven estimation of voxel-wise propagation rates from patient data.

Here, we introduce a differentiable reaction–diffusion framework for tau propagation that overcomes these limitations. By embedding diffusion MRI-derived anisotropy and voxel-wise production dynamics within an end-to-end differentiable simulator, we enable gradient-based estimation of more than one million spatially varying parameters directly from human tau PET data. This approach transforms mechanistic disease modelling into a scalable parameter inference problem and allows subject-specific reconstruction of spatial propagation kinetics across the whole brain.

Applying this framework to tau PET data from the Alzheimer’s Disease Neuroimaging Initiative (ADNI), we reconstruct subject-specific, spatially resolved maps of tau amplification across the human brain. We characterize the spatial organization of the inferred amplification rates using both cortical parcellations and a white-matter tract atlas, revealing systematic regional and tract-specific variation. The inferred amplification rates also show a positive spatial association with amyloid PET, suggesting that they capture aspects of amyloid-associated regional vulnerability. To investigate the molecular basis of the inferred vulnerability landscape, we further integrated the estimated amplification maps with spatial transcriptomic profiles from the Allen Human Brain Atlas (AHBA) [14]. This analysis identified gene expression components associated with the spatial variability of tau amplification through gene set enrichment analysis. Together, these results connect molecular architecture, mechanistic disease dynamics, and predictive modelling of neurodegenerative progression.

## Results

### Differentiable simulation for inferring spatially resolved tau amplification rates

To enable data-driven inference of kinetic parameters governing tau propagation, we first developed a differentiable simulator of pathological tau spreading in the human brain (Fig. 1). To this end, we formulated tau propagation using a reaction–diffusion model with logistic growth, in which the level of pathological tau evolves through diffusion along structural brain connectivity and a local self-catalytic amplification of pathological tau [10]. This model is integrated with an optimization framework, enabling differentiation of the discrepancy between simulated and PET-derived tau levels with respect to model parameters. The dynamics is described by the following equation:

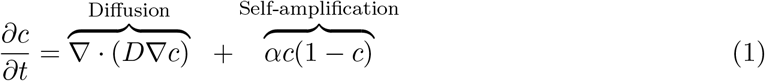

where *c*(**r**, *t*) denotes the normalized level of pathological tau at location **r** = (*x, y, z*) and time *t, D*(**r**) is an anisotropic diffusion tensor derived from diffusion tensor imaging (DTI), and *α*(**r**) represents the local conversion rate from healthy to pathological tau species (See Methods for details). The central goal of this framework is to infer a spatial map of the prion-like self-amplification rate of tau, represented by *α*(**r**), that best explains the observed PET patterns.

**Figure 1:**
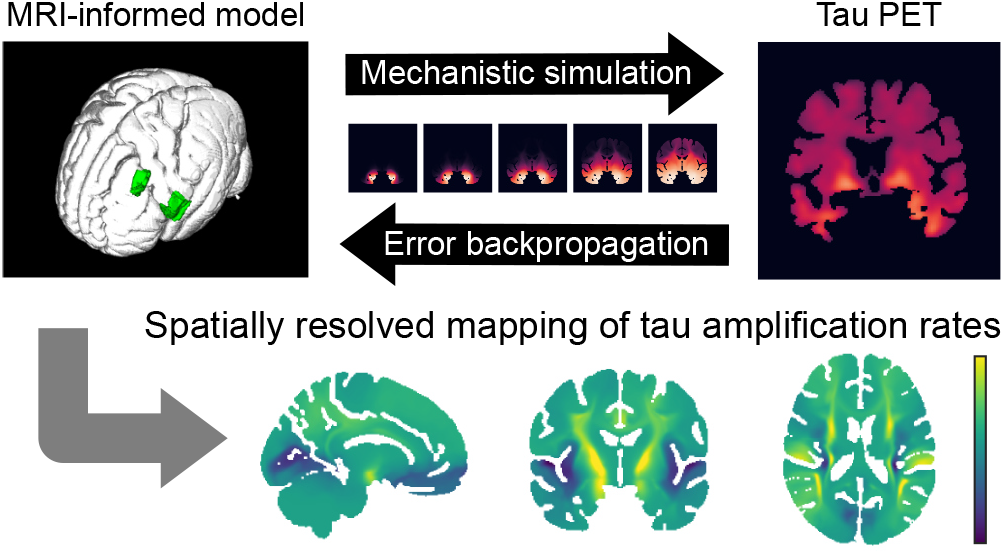
Schematic overview of the proposed framework. A mechanistic reaction–diffusion model informed by MRI data simulates tau propagation initiated from the entorhinal cortex, and the discrepancy with tau PET measurements is minimized via error backpropagation to infer spatially resolved tau amplification rates.

### Fiber-guided diffusion enhances model–PET correspondence

To illustrate the forward model, Fig. 2a shows a representative simulation of tau propagation initiated from the entorhinal cortex, consistent with early Braak-stage pathology. For the simulations shown in Fig. 2, we used a spatially uniform tau amplification rate (*α* = 0.17 [1/year] [15]). In the anisotropic diffusion model, tau spreads preferentially along white-matter fiber tracts, whereas an isotropic control produces more spatially uniform propagation patterns.

**Figure 2:**
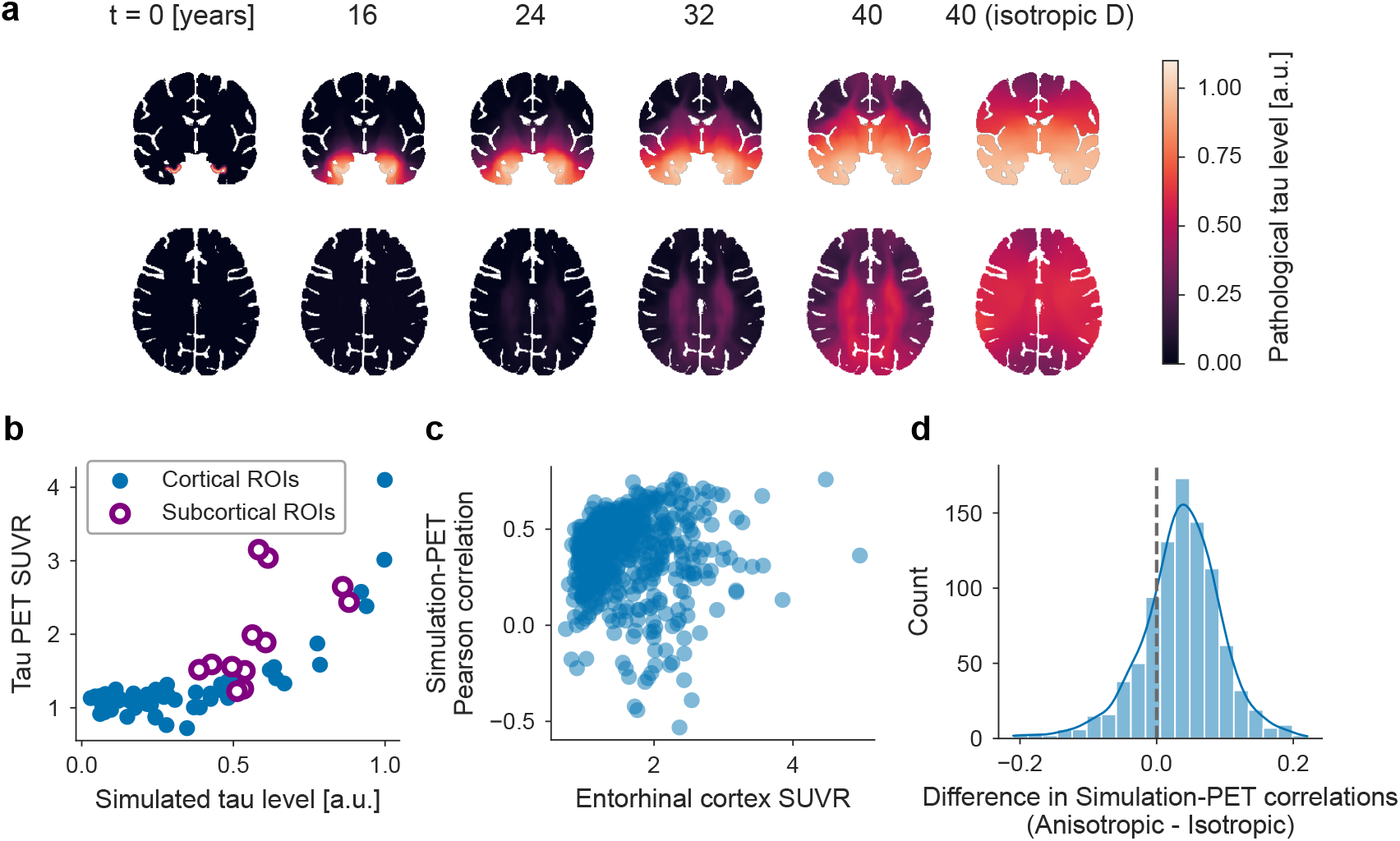
Forward simulation of tau propagation and comparison with tau PET data. (a) Simulated spatiotemporal evolution of pathological tau initiated from the entorhinal cortex, showing propagation along white-matter tracts under anisotropic diffusion, with an isotropic diffusion model shown in the rightmost column as a control. The isotropic model was constructed such that the trace of the diffusion tensor matches that of the anisotropic model. (b) Comparison between simulated tau levels and PET-derived signals across cortical (blue circles) and subcortical (open purple circles) regions using parcellation-based averaging, shown for a representative subject. The Pearson correlation was computed after excluding the entorhinal and temporal pole regions, yielding *r* = 0.67 across all parcels, and *r* = 0.67 when restricted to cortical regions. (c) Relationship between entorhinal cortex tau PET signal and simulation–PET Pearson correlation across individuals. Pearson *r* = 0.035. (d) Distribution of differences in regional correlation between anisotropic and isotropic diffusion models. The vertical dashed line indicates equal performance between the anisotropic and isotropic models. Paired t-test, *p <* 10^*−*8^.

To quantitatively compare simulated tau distributions with PET measurements, we adopted a coarse-grained similarity metric based on cortical and subcortical parcellation (Fig. 2b), shown here as a representative plot from a single individual. Specifically, the brain was partitioned according to the Desikan–Killiany (DK) atlas, and parcel-wise mean PET intensities were compared with the corresponding mean simulated tau levels. The PET measurements were obtained from the processed and quantified tau-PET dataset provided by the ADNI PET Core. The Pearson correlation across all parcels was then used as a similarity score, excluding the entorhinal cortex and temporal pole, which consistently exhibit high signal levels. In the representative example shown in Fig. 2b, this yielded a Pearson correlation of *r* = 0.67. Compared with voxel-wise measures, the parcellation-based measure bypasses the need for precise spatial alignment between simulated and measured signal distributions. Because it relies on correlation-based comparison, it also eliminates the need for additional signal normalization. Across individuals, the resulting simulation–PET correlation showed only a weak association with entorhinal tau levels (Pearson *r* = 0.035; Fig. 2c), suggesting that model performance is not strongly driven by disease stage.

We quantitatively compared the anisotropic diffusion model with an isotropic control to assess whether incorporating DTI-derived structural information improves the reproduction of tau PET patterns. We sampled more than 900 individuals from the dataset and evaluated the simulation–PET similarity based on parcellation for each subject. As shown in Fig. 2d, the anisotropic model consistently achieved higher regional correlation scores (paired t-test, *p <* 10^*−*8^). This indicates that incorporating fiber-guided diffusion substantially improves the ability of the model to reproduce the spatial distribution of tau pathology observed in vivo, even under spatially uniform tau amplification rate.

### Data-driven inference reveals structured heterogeneity in tau amplification

Next, we present brain maps of tau amplification rates inferred to reproduce the PET data (Fig. 3). To this end, we first describe the training setup. Gradient-based learning requires the specification of hyperparameters such as the learning rate and the number of epochs. However, because our framework estimates parameters individually for each subject, conventional training–test splits for cross-validation are not applicable. Instead, hyperparameters were determined using a subset of 100 individuals, and the final training was performed only once on 166 individuals selected from the remaining 827 subjects with the highest tau PET signal in the entorhinal cortex (See Methods for details).

**Figure 3:**
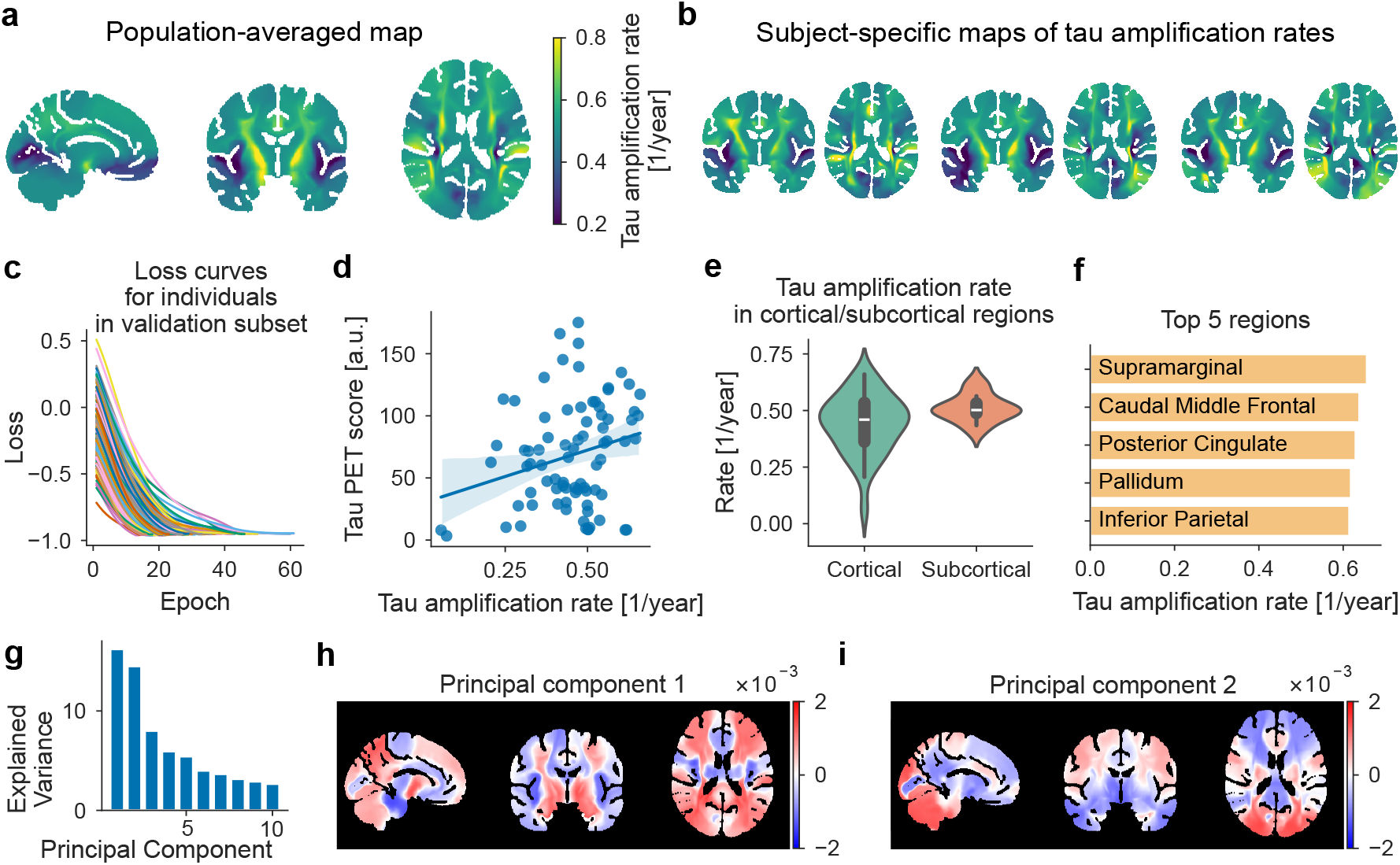
Data-driven reconstruction of spatially heterogeneous tau amplification rates. (a) Population-averaged map of the inferred tau amplification rates. (b) Subject-specific amplification maps. Coronal and axial slices are shown for three representative individuals. The color scale is the same as in (a). (c) Loss curves for individuals in the validation subset. (d) Relationship between regional tau PET scores and inferred amplification rates, both averaged across individuals. Data points represent cortical and subcortical brain regions. Linear regression line with 95% confidence intervals is shown. Pearson *r* = 0.25. (e) Distribution of amplification rates across cortical and subcortical regions. (f) Bar plot of brain regions exhibiting the highest amplification rates. (g) Explained variance of principal components derived from individual amplification maps. (h,i) Spatial patterns of the first and second principal components, capturing dominant modes of variability.

Fig. 3a shows the population-averaged amplification map, which reveals a structured spatial pattern across the brain rather than a uniform distribution. At the individual level, the inferred maps share broadly similar spatial patterns, consistent with common patterns of tau propagation, while still exhibiting substantial inter-individual variability, as illustrated by representative subject-specific maps (Fig. 3b). To quantify this variability, we computed the variance-to-mean ratio of amplification rates across individuals, which revealed high variability in the transverse temporal, parahippocampal, and insular regions. To confirm the stability of the optimization, we examined the loss trajectories during training. The loss consistently decreased and converged across individuals (Fig. 3c), indicating that gradient-based learning successfully optimized voxel-wise parameters.

We next examined the relationship between the inferred amplification rates and regional tau PET signals after baseline removal (see Methods). Both quantities were evaluated at the population-average level. The correspondence is limited (Pearson *r* = 0.25; Fig. 3d), indicating that the inferred amplification reflects latent kinetic properties that are not trivially explained by the observed tau burden.

Regional analysis further revealed systematic differences between brain compartments. Amplification rates exhibited distinct distributions between cortical and subcortical regions (Fig. 3e). In particular, cortical regions showed high variability, indicating heterogeneity in cortical vulnerability. Then we examined regional profiles of high amplification (Fig. 3f). Several regions, including the supramarginal and caudal middle frontal cortices, exhibited elevated amplification rates. Notably, these regions do not show particularly high tau burden, suggesting that amplification rate captures latent regional susceptibility that is not directly reflected in observed tau accumulation. We further examined the inferred amplification rates across major white-matter tracts (Supplementary Fig. 2). Higher amplification rates tended to occur in tracts farther from the entorhinal cortex, including frontoparietal association pathways.

To characterize inter-individual variability, we performed principal component analysis on the inferred individual maps. The variance explained by each component is shown in Fig. 3g, with the first and second components capturing a large fraction of variability. The spatial patterns of the leading components (Fig. 3h,i) visualize dominant modes of variability. In the coronal view, PC1 reflects a medial–lateral contrast, whereas PC2 captures a superior–inferior gradient. These results collectively indicate that tau amplification exhibits structured spatial organization and substantial inter-individual variation that is distinct from observed PET distributions.

### Spatial association of tau amplification rates with amyloid pathology

Because the inferred amplification rates capture spatial features that are distinct from the observed tau burden, we next examined their relationship with amyloid pathology. For each individual, we computed the spatial correlation between the inferred tau amplification rates and amyloid PET signals across brain regions (Fig. 4a). Positive correlations were observed in most individuals. Moreover, the amplification rate-amyloid PET correlation was significantly higher than the tau PET-amyloid PET correlation used as a baseline. These results suggest that the inferred amplification rates capture spatial features associated with amyloid pathology beyond those directly reflected in the tau PET distribution. We also examined the regional contributions to the correlation coefficients and found heterogeneous positive and negative contributions across brain regions (Fig. 4b).

**Figure 4:**
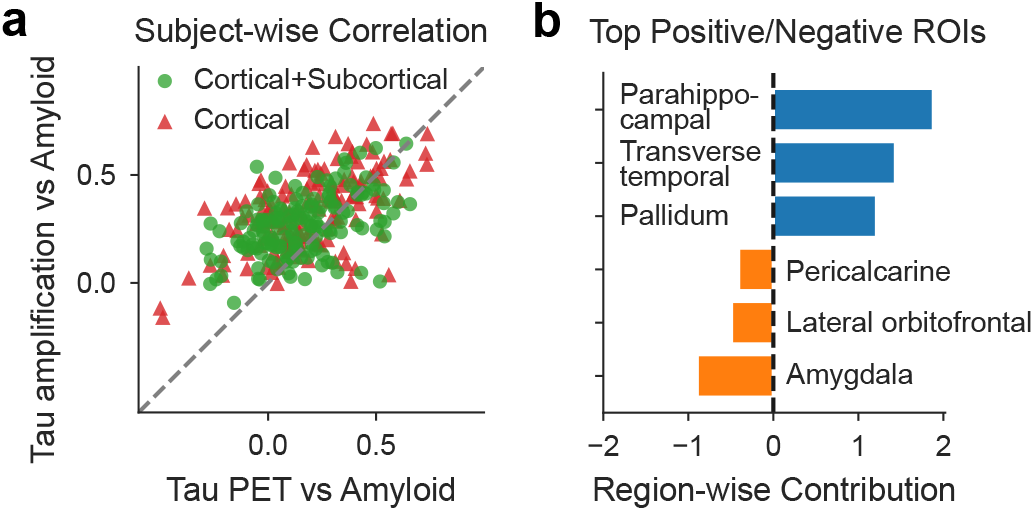
Spatial association of tau amplification rates with amyloid PET signals. (a) Subject-wise spatial correlations with amyloid PET. For each individual, the spatial correlation between inferred tau amplification rates and amyloid PET is compared with the tau PET–amyloid PET correlation as a baseline. Results are shown using cortical and subcortical regions together (green circles) and cortical regions only (red triangles). The dashed line indicates equal correlation with amyloid PET. The amplification rate–amyloid PET correlation was significantly higher than the tau PET–amyloid PET correlation in both analyses (paired t-test, (*p <* 10^*−*10^) for cortical and subcortical regions; (*p <* 10^*−*10^) for cortical regions only). (b) Brain regions showing the largest positive and negative contributions to the spatial correlation between tau amplification rates and amyloid PET.

### Gene expression patterns predict spatial variation in tau amplification

We next sought to investigate the molecular basis of regional variation in tau amplification. To this end, we leveraged spatial gene expression data from the AHBA, which provides transcriptomic profiles across the human brain (Fig. 5).

**Figure 5:**
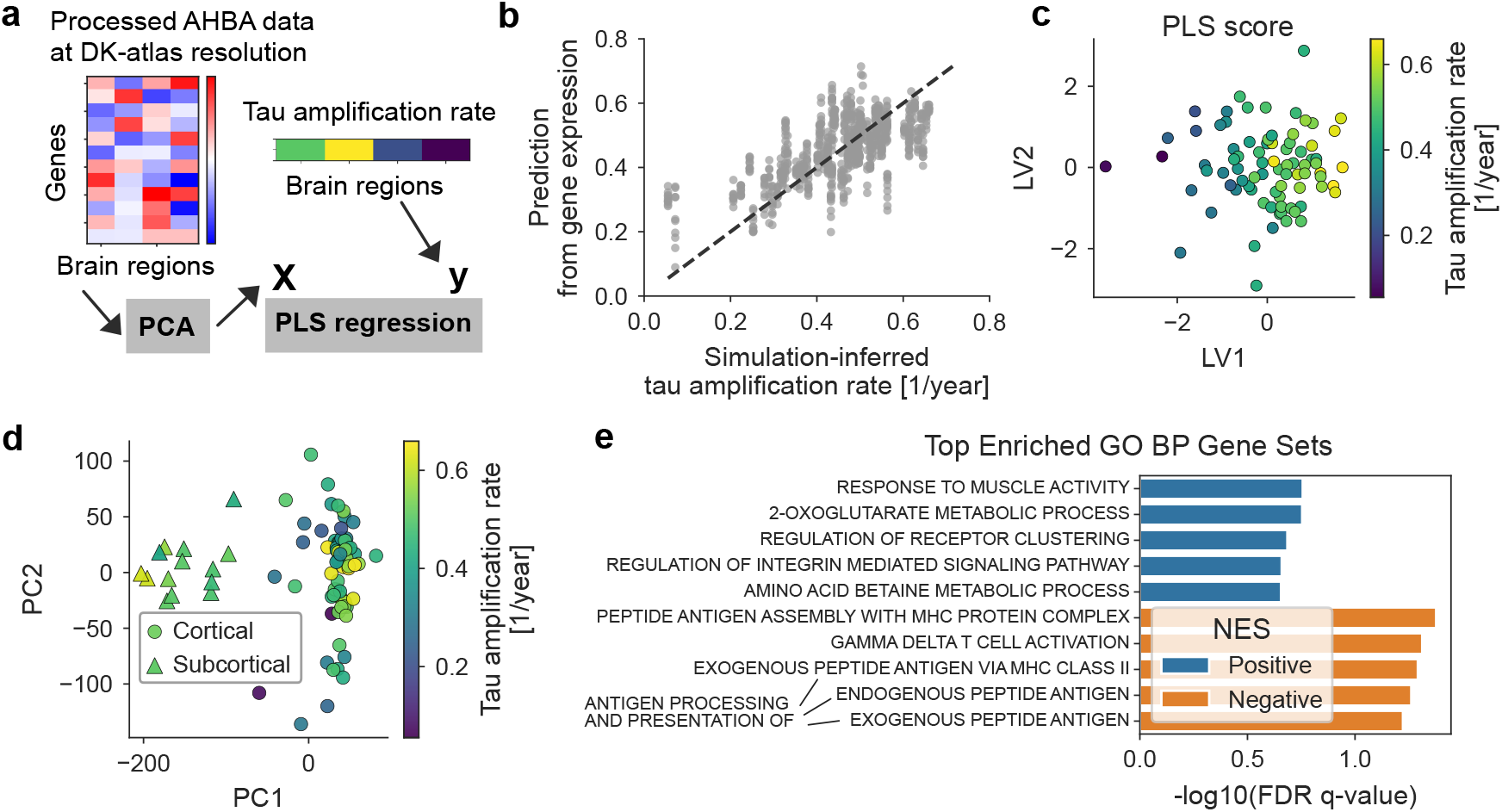
Transcriptomic correlates of spatial variation in tau amplification rates. (a) Workflow of the analysis integrating AHBA gene expression data with inferred tau amplification rates using partial least squares (PLS) regression. (b) Cross-validated prediction of amplification rates from gene expression. Cross-validation with random partitioning was repeated 10 times, yielding 10 predicted values for each brain region, all of which are shown. (c) Low-dimensional representation of gene expression defined by PLS, where each point corresponds to a brain region and is colored according to its tau amplification rate. (d) Low-dimensional representation of gene expression defined by PCA. Circles and triangles denote cortical and subcortical regions, respectively. (e) Gene set enrichment analysis (GSEA) of the leading PLS component (LV1), showing positively and negatively enriched Gene Ontology biological process terms.

We first examined the relationship between the inferred tau amplification rates and canonical Alzheimer’s disease-related genes, MAPT and APOE. The expression levels of these genes have been shown to correlate with regional tau PET signals [16, 17, 18]. However, their association with the amplification rates was substantially weaker (Supplementary Fig. 3), consistent with the distinct spatial patterns of tau burden and amplification dynamics. We then performed supervised dimension reduction using partial least squares (PLS) regression [19, 8, 20], to identify multivariate gene expression patterns associated with tau amplification (Fig. 5a). This analysis extracts latent components in the high-dimensional gene expression space that maximally covary with the amplification rates. Under cross validation, predictions of amplification rates based on gene expression were significantly better than those obtained from spatially autocorrelated surrogate data (*p* = 0.0012) [21], indicating that the observed associations cannot be explained by spatial smoothness alone (Fig. 5b) (See Methods and Supplementary Fig. 4). Fig. 5c shows the PLS-defined low-dimensional representation of gene expression across brain regions, colored by the corresponding amplification rates. Regions with high and low amplification rates were well separated in this space. In our setting, only the first component (LV1) was correlated with the tau amplification rates; therefore, we focus on the analysis of LV1. Before performing enrichment analysis on LV1, we also examined the low-dimensional representation obtained by principal component analysis (PCA). Although cortical and subcortical regions were clearly separated in this space as shown in Fig. 5d, separation by amplification rates was substantially weaker.

To interpret the extracted gene expression patterns by PLS analysis, we performed gene set enrichment analysis (GSEA) [22] on the leading component (LV1), which captures the dominant axis of variation in gene expression associated with amplification rates. Fig. 5e shows the top 5 positively and top 5 negatively enriched gene sets from the Gene Ontology Biological Process (GO BP) database. Gene sets positively associated with tau amplification were primarily related to metabolic processes and cellular signaling, whereas gene sets negatively associated with tau amplification were dominated by immune-related pathways, including antigen processing and T cell activation. These results indicate that the spatial patterns of tau amplification rates inferred from PET data are associated with specific molecular programs, supporting the biological relevance of the inferred kinetic landscape.

Finally, we examined whether subject-specific amplification rates exhibit systematic differences across demographic and genetic subgroups, focusing on sex and APOE genotype (Supplementary Fig. 5). No marked differences were observed between male and female subjects or between APOE *ε*4 carriers and non-carriers. This may reflect the cohort selection, which was restricted to individuals with pronounced tau accumulation in the entorhinal cortex, with most subjects being amyloid-positive (144 positive, 13 negative, and 9 missing), potentially masking subgroup effects.

## Discussion

In this study, we established a framework for inferring spatially resolved tau amplification rates directly from subject-specific PET data using a differentiable reaction–diffusion model. In the forward analysis (Fig. 2), we showed that DTI-based anisotropic diffusion improves the reproduction of in vivo tau patterns compared with isotropic models, supporting the role of fiber-guided propagation at the whole-brain scale. In the inverse setting (Fig. 3), the inferred amplification maps revealed a structured and non-uniform spatial organization that is consistent across individuals yet exhibits substantial inter-individual variability. Notably, amplification rates showed only limited correspondence with regional tau PET burden, indicating that they capture latent kinetic properties rather than simply reflecting observed pathology. Analysis of white-matter tracts further revealed elevated amplification rates in several frontoparietal pathways relatively distant from the entorhinal cortex (Supplementary Fig. 2). In the current reaction-diffusion model (Eq. 1), the propagation velocity of pathology (*v*) is scaled with the amplification rate (*α*) as 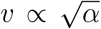. Thus, tract-specific variation in the inferred amplification rates may reflect heterogeneity in effective tau propagation along axonal pathways, although the underlying neurobiological mechanisms remain unclear.

The spatial organization of the inferred amplification rates was also associated with amyloid pathology (Fig. 4). Across individuals, the inferred amplification rates showed a stronger spatial association with amyloid PET than tau PET did, despite amyloid information not being used in the inference. Previous studies have demonstrated a close relationship between amyloid burden and the spatial distribution of tau pathology, including the extension of tau into neocortical regions [23, 2]. Our findings extend this relationship from observed tau burden to the inferred kinetics of tau amplification, providing a mechanistic perspective on the spatial coupling between amyloid and tau pathology.

Our transcriptomic analysis provides additional insight into the biological basis of the inferred amplification landscape (Fig. 5). Gene expression patterns associated with amplification rates differed from those linked to tau distribution, and enrichment analysis highlighted metabolic and immune-related pathways. One possible interpretation is that gene expression programs positively associated with tau amplification reflect regions with higher neuronal activity, which may be more susceptible to amyloid accumulation and thereby facilitate subsequent tau propagation [24, 25, 26]. In contrast, gene sets negatively associated with tau amplification were primarily related to immune processes, potentially reflecting molecular programs linked to resilience or protective responses [27]. Importantly, the AHBA data are derived from neurologically normal donors. Therefore, these associations should not be interpreted as disease-induced transcriptional responses, but rather as baseline regional properties that shape differential vulnerability and resilience to pathological processes.

Several limitations of the present framework should be noted. First, the diffusion process was modeled using DTI-derived tensors that treat all white-matter tracts equivalently, whereas in reality, axonal pathways may differ in their capacity to support trans-synaptic propagation. This simplification may contribute to discrepancies between simulated and observed propagation patterns along specific tracts. Second, while we focused on estimating the spatial heterogeneity of the amplification rate, the diffusion coefficient was fixed. Extending the framework to jointly infer diffusion parameters would be particularly informative, as recent studies suggest that tau propagation may involve multiple transport mechanisms, including both synaptic transmission and extracellular or vesicle-mediated processes [28]. In this context, isotropic diffusion may partially capture non-synaptic spreading, and disentangling these contributions through parameter inference could provide a more complete characterization of propagation modes and their regional heterogeneity. Third, the absolute scale of the inferred tau amplification rate is not directly identifiable. Although the assumed timescale (*α ∈* [0, 1] year^*−*1^) yields reasonable values, it remains a modeling choice, and thus only the relative spatial variation of amplification rates should be interpreted. Incorporating longitudinal PET data would enable estimation of its absolute scale. Fourth, most of our analyses relied on spatial coarse-graining based on brain parcellation, which does not fully exploit the high spatial resolution of inferred maps. Recent efforts have sought to reconstruct spatially dense, neuroimaging-compatible transcriptomic maps from the AHBA data [29], and integrating such approaches into our pipeline would enable a more fine-grained characterization of regional heterogeneity.

Finally, the proposed inverse mapping framework opens several avenues for future applications. With the advent of disease-modifying therapies targeting pathological proteins, such as anti-amyloid antibodies, longitudinal imaging datasets acquired under therapeutic intervention will become increasingly available. Applying the present approach to such data could enable quantitative assessment of how treatments alter local propagation kinetics, providing a mechanistic readout beyond conventional measures of pathology burden. Moreover, while we focused on transcriptomic correlates, the inferred amplification maps can be readily compared with other spatial brain datasets, including PET markers of neuroinflammation such as microglial activation. Such multimodal integration may help disentangle the biological drivers of propagation dynamics. The framework is also extendable to other neurodegenerative diseases as imaging probes become available, including emerging PET ligands for alpha-synuclein. We anticipate that the inverse mapping method will contribute to a mechanistic and predictive understanding of neurodegenerative disease progression.

## Methods

### Model

As described in Eq. (1) of the main text, we model the dynamics of the abnormal protein concentration *c*(**r**, *t*), where **r** = (*x, y, z*) denotes the spatial location in the brain, using the following reaction-diffusion equation [10, 11]:

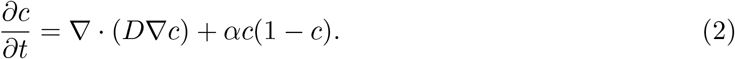

Here, the conversion of normal proteins into abnormal ones is represented by a logistic reaction term with rate *α*(r), while the propagation of abnormal proteins is modeled by an anisotropic and spatially heterogeneous diffusion tensor *D*(r) reflecting the underlying white matter architecture. As detailed below, the diffusion tensor *D*(r) is constructed from diffusion tensor imaging (DTI) data.

At the brain surface, we imposed the no-flux boundary condition, **n** *· D∇c* = 0, where **n** is the unit vector orthogonal to the brain surface. To simulate the complex brain geometry, various approaches such as the finite element method have been used. In this study, we adopted the phase-field method [30], which supports structured grids, reduces memory usage, and simplifies the design of inverse problems. Introducing an auxiliary variable *ϕ* (r), which satisfies *ϕ* = 1 inside the brain and *ϕ* = 0 outside, the reaction-diffusion equation becomes:

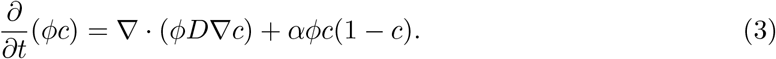

### Numerical scheme

For numerical integration of Eq. 3, we first computed the numerical fluxes *ϕD∇c* on staggered grids by central differencing and then approximated the divergence via flux differencing. The diffusion term was combined with the reaction term *αϕc*(1 *− c*), and the temporal update was carried out by the explicit Euler scheme, following previous studies on simulating phase-field models [31, 32]. The simulation grid contained 182 *×* 218 *×* 182 nodes, mirroring the voxel dimensions of the MNI standard brain template. Space and time were discretized with Δ*x* = 1 [mm] and Δ*t* = 10^*−*4^ [years], respectively. We note that each voxel represents a volume of 1mm^3^, which, in vivo, typically encompasses on the order of 10^5^ neurons and glial cells. At this resolution, the diffusion term implicitly captures sub- and supra-cellular transport mechanisms, such as axonal transport, vesicle secretion, and cell migration, that operate below the voxel scale.

We implemented the numerical simulation using JAX (version 0.4.26), a Python library that supports automatic differentiation and GPU acceleration [33]. As a result, derivatives of a loss function required for gradient-based parameter optimization can be easily computed via automatic differentiation. All computations were performed on an NVIDIA RTX 6000 Ada GPU. The implementation of the differentiable simulation, designed to minimize dependencies on external libraries, will be made publicly available via a GitHub repository.

### Tau PET

Tau PET data used in the preparation of this article were obtained from the Alzheimer’s Disease Neuroimaging Initiative (ADNI) database (adni.loni.usc.edu). The ADNI was launched in 2003 as a public-private partnership, led by Principal Investigator Michael W. Weiner, MD. The primary goal of ADNI has been to test whether serial magnetic resonance imaging (MRI), positron emission tomography (PET), other biological markers, and clinical and neuropsychological assessment can be combined to measure the progression of mild cognitive impairment (MCI) and early Alzheimer’s disease (AD).

### Neuroimage analysis and model parametrization

The brain geometry was defined based on a population-averaged structural magnetic resonance imaging (MRI) template. We used the MNI152 standard-space T1-weighted template (MNI152 nonlinear 6th generation asymmetric), distributed with FSL (version 6.0.7.7) [34], and processed it using the `recon-all` pipeline in FreeSurfer (version 7.4.1) [35]. The resulting `aparc+aseg` volume was used to obtain a comprehensive parcellation of subcortical and cortical gray matter regions, combining automated subcortical segmentation with cortical parcellation based on the Desikan-Killiany atlas. We also extracted three binary masks: brain parenchyma, white matter, and gray matter. The brain parenchyma mask was evolved in time using the Allen-Cahn equation to smooth the boundary at the brain surface, and the resulting field was used as the phase-field auxiliary variable *ϕ*(**r**). The masks of white and gray matters are used to set the diffusion tensor field in simulation as described below.

For parametrization of the diffusion tensor field, we adopted the population-averaged diffusion tensor image (DTI) constructed from 1065 subjects in the Human Connectome Project (HCP) data, which is also provided with the FSL software [34, 36]. From this data, the principal eigenvector ***γ*** was computed at each voxel, and the diffusion tensor for the simulation was defined as:

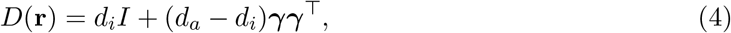

following [10]. In white matter, we set *d*_*i*_ =1 mm^2^/year and *d*_*a*_ =100 mm^2^/year, while in gray matter, we used *d*_*i*_ = *d*_*a*_ =10 mm^2^/year.

### Inference of tau amplification rates from PET

The voxel-wise tau amplification rates, *α*(**r**), were updated based on the gradient of the discrepancy between simulated and measured tau distributions, i.e.,

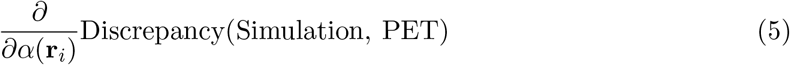

where **r**_*i*_ is the location of the *i*-th voxel. As a criterion to quantify this discrepancy, we used the negated Pearson correlation between simulated and measured tau levels across brain regions defined by parcellation. Specifically, the discrepancy quantification used only cortical regions, excluding subcortical regions. In addition, the entorhinal cortex and temporal pole were omitted, as these regions consistently exhibit high tau signals and could otherwise lead to an overestimation of model goodness of fit.

The target tau PET signals used for training were constructed from the ADNI UC Berkeley tau PVC dataset (“16Dec2024” version) as follows. For each individual with flortaucipir (FTP) scans, we selected the most recent acquisition and excluded subjects with missing values in the entorhinal cortex, resulting in 927 individuals. These were split into a development subset (100 individuals) and a validation subset (827 individuals) using stratified sampling based on tau PET levels in the entorhinal cortex. Within each subset, individuals were further divided into five groups according to entorhinal tau levels. Using the lowest 20% group as a baseline distribution, we computed one-sample t-statistics at each brain region for the top 20% individuals with the highest tau accumulation as the target tau PET signals for training. This procedure, following previous studies, was designed to reduce the influence of basal signal distributions unrelated to neurodegenerative pathology. As a result, we obtained 20 and 166 individuals in the development and validation subsets, respectively, as target tau distributions. To determine algorithmic hyperparameters for the optimization, gradient-based learning was performed on the 20 individuals in the development subset under various settings (Supplementary Fig. 1). Based on these experiments, we selected a configuration that achieved sufficient optimization while avoiding overfitting and oscillations of the objective function: 60 epochs in total, cosine annealing starting from epoch 40, and early stopping when the Pearson correlation exceeded

0.95. The initial learning rate was set to 1, with gradient clipping applied to ensure that the L2 norm of the gradient did not exceed 10. The simulation time was 20 years. The tau amplification rate *α* was constrained to lie within the range [0, 1] [37, 10, 15], and was initially set to a spatially uniform value of 0.5. Under these settings, spatially resolved maps of tau amplification rates were optimized for the 166 individuals in the validation subset.

### White-matter tract analysis of tau amplification rates

For tract-level analysis of the inferred tau amplification rates, we used the HCP-derived XTRACT white-matter tract atlas distributed with FSL [38]. For each tract, the tau amplification rate was calculated by averaging the voxel-wise inferred rates within the corresponding tract mask. To examine whether amplification rates vary with spatial location, we calculated the Euclidean distance between the centroid of each white-matter tract mask and that of the entorhinal cortex mask. Distances were calculated separately for the left and right hemispheres and then averaged.

### Association of tau amplification rates with amyloid PET

For each individual, we selected the amyloid PET scan closest in time to the tau PET scan from the ADNI UC Berkeley amyloid PET dataset (“03Jun2025” version). If the nearest amyloid PET scan was more than five years apart from the tau PET scan, the subject was excluded from this analysis. To assess the spatial association between tau amplification and amyloid pathology, we computed the Pearson correlation between inferred amplification rates and amyloid PET signals across brain regions for each individual. We then decomposed this correlation into regional contributions to identify which areas contributed strongly to the observed association. For amplification rate (*a*_*i*_) and amyloid PET signal (*m*_*i*_) in the *i*-th region, the Pearson correlation can be written as Σ_*i*_ *s*_*i*_, where the contribution score of the *i*-th region is defined as

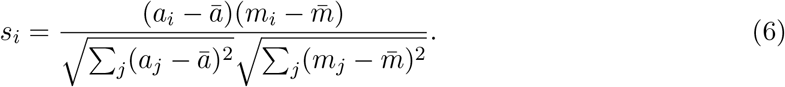

These region-wise contributions were calculated for each individual and then averaged across individuals.

### Cohort characterization and subgroup analyses

Demographic, genetic, and amyloid status information for these individuals in the validation subset was obtained from ADNI study data files. These variables were used for subgroup comparisons of subject-specific amplification maps and for characterizing the composition of the analyzed cohort. For amyloid status, we identified the amyloid PET scan date closest to the tau PET scan date for each individual and used the corresponding amyloid status. If the nearest amyloid PET scan was more than five years apart from the tau PET scan, the amyloid status was not used.

### Supervised dimension reduction of gene expression matrix

To characterize spatial patterns of transcriptional profiles in the human brain, we used the Allen Human Brain Atlas (AHBA) dataset [14]. We preprocessed the AHBA data and mapped them to the brain regions of interest using the abagen toolbox [39, 40]. As a result, consistent with the tau PET signals, regional gene expression profiles were obtained for 80 cortical and subcortical gray matter regions. The regional expression profiles were averaged across donors, and missing values were imputed using the `centroid` option. The resulting gene expression matrix was reduced to 30 dimensions using principal component analysis (PCA). We then performed supervised dimension reduction using partial least squares (PLS) regression, with the population-averaged tau amplification rates inferred from simulations as the response variable. In practice, we used the `PLSRegression` implementation in the scikit-learn library (version 1.6.1). For cross-validation, we employed structured partitioning to avoid information leakage arising from the similarity of gene expression patterns between left–right homologous brain regions. Specifically, corresponding regions were always assigned to the same data subset. To evaluate the statistical significance of the goodness of fit, we used *R*^2^ regression score and surrogate data that preserve spatial autocorrelation. The surrogate datasets (10^4^ samples) were generated by the BrainSMASH method [21]. To model the spatial autocorrelation, we used the Euclidean distance between the centroids of cortical and subcortical regions, which is supported in part by a recent finding of whole-brain molecular gradients in the adult brain [20]. In the analyses shown in Fig. 5c and thereafter, we present the PLS results obtained using all data points under the same settings. Finally, for gene set enrichment analysis (GSEA), gene-wise contributions to the first PLS component (LV1) were computed as follows. Let the full gene expression matrix and its PCA representation be related as

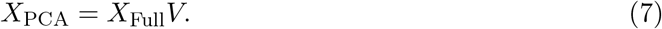

Then let *D*^*−*1^ be the diagonal scaling matrix applied to the inputs in PLSR, and *R* the transformation matrix of PLSR. The PLSR score matrix *T* can be expressed as

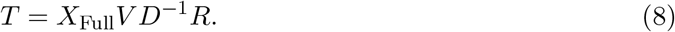

We therefore computed the first column of *V D*^*−*1^*R* for the contribution of each gene to LV1.

### Enrichment analysis

Gene ranking was constructed based on the contributions of each gene to the PLS score of the first component (LV1). Gene set enrichment analysis (GSEA) was performed using the GSEA software (version 4.4.0, Broad Institute) in preranked mode with GO Biological Process gene sets (c5.go.bp.v2026.1.Hs.symbols.gmt). Statistical significance was assessed based on FDR q-values.

## Supplementary information

Supplementary information, including Supplementary Figures 1–5, accompanies this manuscript.

## Acknowledgements

This research was supported by the Moonshot R&D–MILLENNIA Program (JPMJMS2024-9 to Y.K. and H.N.) from Japan Science and Technology Agency (JST) and JST PRESTO (JPMJPR25K3 to Y.K.).

Data collection and sharing for the Alzheimer’s Disease Neuroimaging Initiative (ADNI) is funded by the National Institute on Aging (National Institutes of Health Grant U19AG024904). The grantee organization is the Northern California Institute for Research and Education. In the past, ADNI has also received funding from the National Institute of Biomedical Imaging and Bioengineering, the Canadian Institutes of Health Research, and private sector contributions through the Foundation for the National Institutes of Health (FNIH) including generous contributions from the following: AbbVie, Alzheimer’s Association; Alzheimer’s Drug Discovery Foundation; Araclon Biotech; BioClinica, Inc.; Biogen; Bristol-Myers Squibb Company; CereSpir, Inc.; Cogstate; Eisai Inc.; Elan Pharmaceuticals, Inc.; Eli Lilly and Company; EuroImmun; F. Hoffmann-La Roche Ltd and its affiliated company Genentech, Inc.; Fujirebio; GE Healthcare; IXICO Ltd.; Janssen Alzheimer Immunotherapy Research & Development, LLC.; Johnson & Johnson Pharmaceutical Research & Development LLC.; Lumosity; Lundbeck; Merck & Co., Inc.; Meso Scale Diagnostics, LLC.; NeuroRx Research; Neurotrack Technologies; Novartis Pharmaceuticals Corporation; Pfizer Inc.; Piramal Imaging; Servier; Takeda Pharmaceutical Company; and Transition Therapeutics.

## Author contributions

Y.K and H.N designed the research. Y.K. implemented the model and analyzed the data. Y.K. and H.N wrote the manuscript.

## Data availability

Data from ADNI can be requested via the LONI website (https://adni.loni.usc.edu/).

## Code availability

The source code for simulation and data analysis is publicly available at https://github.com/kondo-y/differentiable-tau-propagation.

