## Supplementary Figures 1-5 for "Spatially resolved mapping of tau amplification rates via differentiable simulation of prion-like propagation"

[*]Data used in preparation of this article were obtained from the Alzheimer's Disease Neuroimaging Initiative (ADNI) database (adni.loni.usc.edu). As such, ADNI investigators contributed to the design and implementation of the ADNI database and/or provided data but did not participate in the analysis or writing of this report. A complete listing of ADNI investigators can be found at https://adni.loni.usc.edu/wp-content/uploads/how_to_apply/ADNI_Acknowledgement_List.pdf.

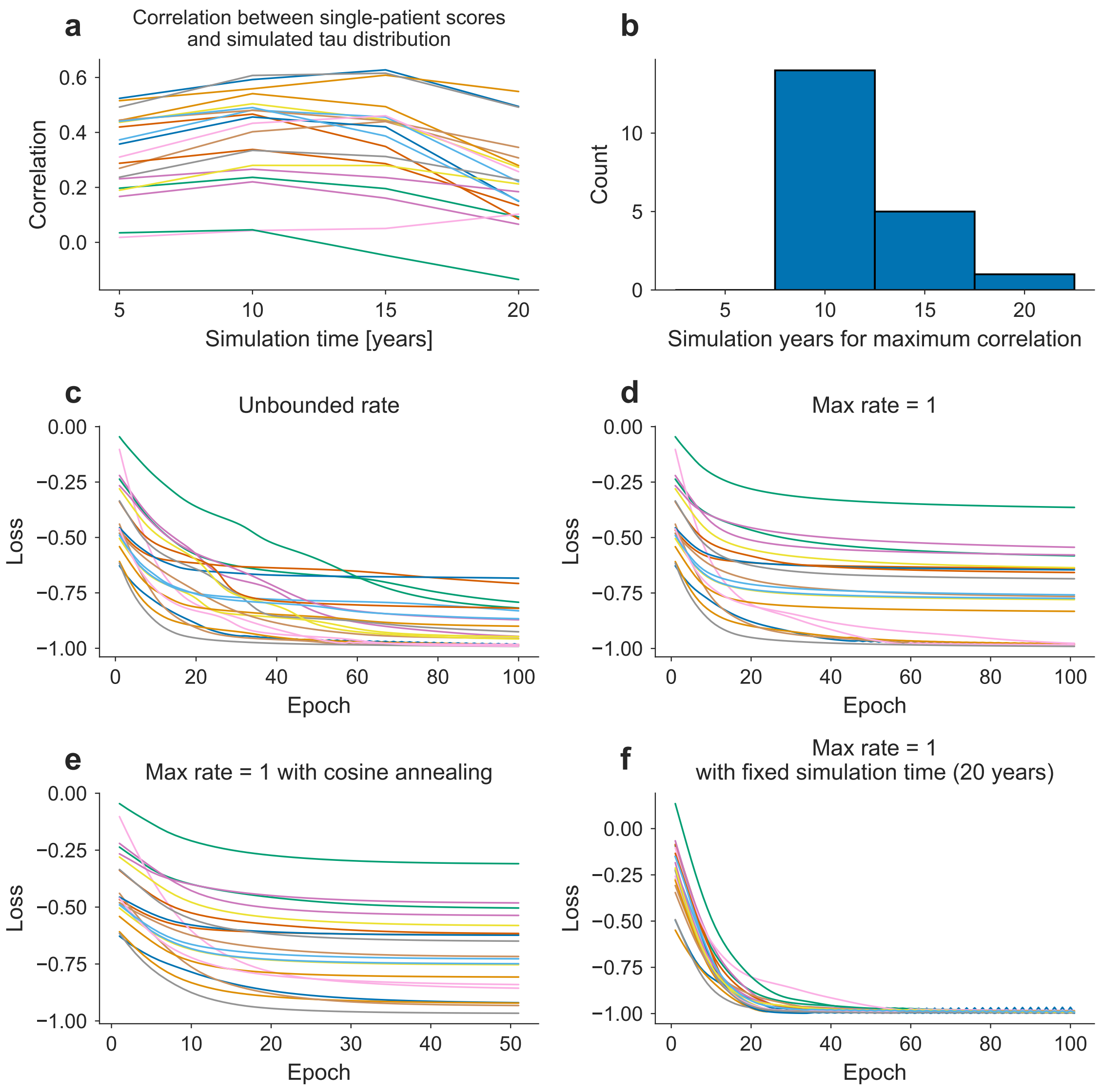


Supplementary Fig. 1: Hyperparameter tuning of gradient-based learning using a development subset. Inference of tau amplification rates was performed on individual patients in a development subset to determine hyperparameters. (a) Evolution of the Pearson correlation between simulated tau distributions and PET-derived tau scores, starting from a uniform tau amplification rate (0.5 [1/year]). Each line represents an individual patient. (b) Histogram of the simulation time at which the simulation–data correlation is maximized. (c) Learning curves for each patient when the simulation time is fixed to the value that maximizes the correlation under the uniform-rate initialization of tau amplification rate. (d) Same as (c), but with the tau amplification rate constrained to the range [0, 1]. (e) Same as (d), with cosine annealing applied to the learning rate. The oscillations in the loss disappear. (f) Same as (d), but with the simulation time fixed to 20 years for all patients.

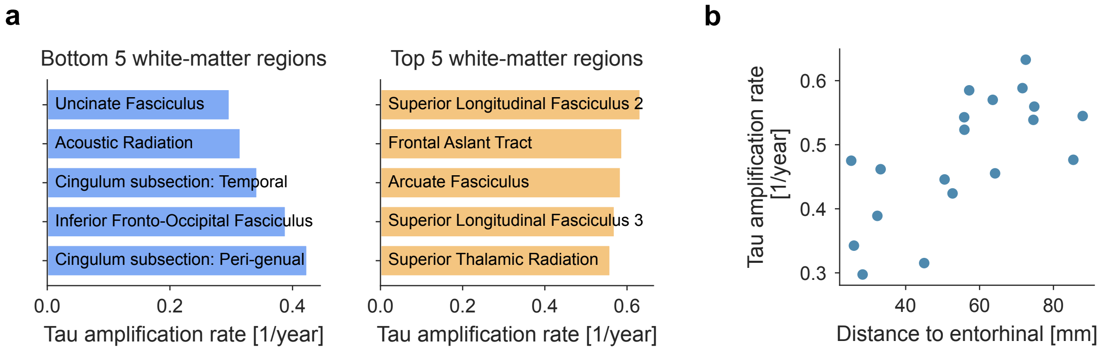


Supplementary Fig. 2: Regional variation in tau amplification rates across white-matter tracts. (a) The five white-matter regions with the lowest (left) and highest (right) estimated tau amplification rates. White-matter tracts were defined using the HCP-derived XTRACT atlas. (b) Relationship between the tau amplification rate and distance from the entorhinal cortex across white-matter regions. Each point represents a white-matter region.

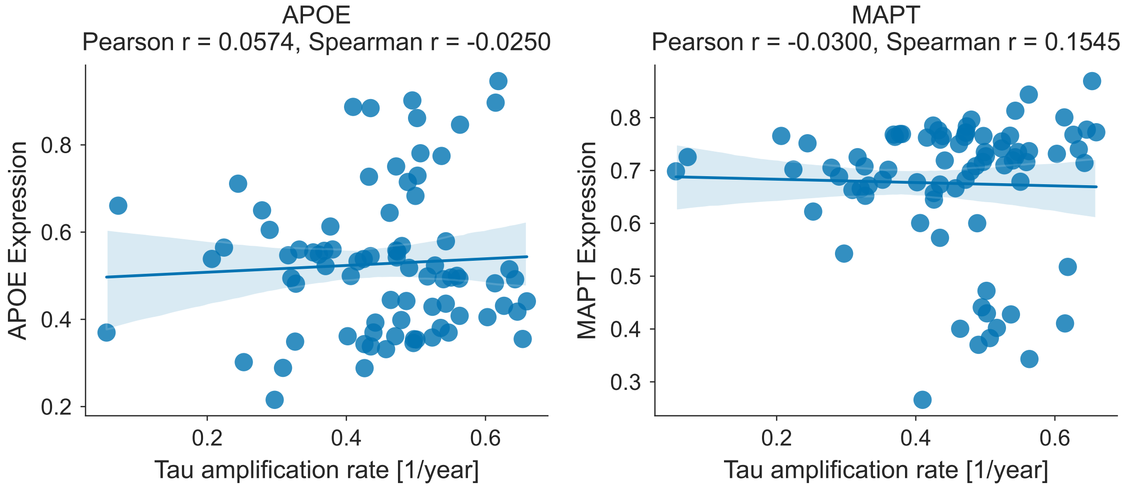


Supplementary Fig. 3: Correlation analysis between simulation-inferred tau amplification rates and expression levels of APOE (left) and MAPT (right) across brain regions. Data points represent brain regions from the Desikan-Killiany atlas and FreeSurfer subcortical segmentation. Linear regression lines with 95% confidence intervals are shown. Pearson and Spearman correlation coefficients are indicated in each panel. Gene expression data from the Allen Human Brain Atlas was processed using the abagen toolbox.

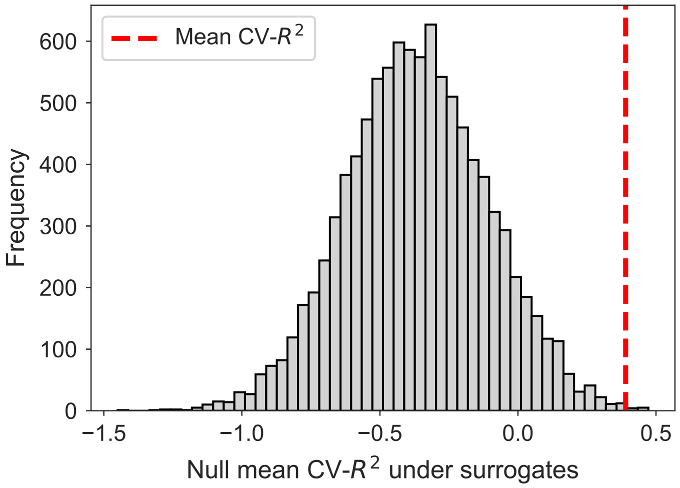


Supplementary Fig. 4: Histogram of cross-validated $R^2$ under surrogate data keeping spatial auto-covariance. The surrogate data were generated by using BrainSMASH method. For each of the 10,000 surrogate datasets, we used the mean $R^2$ obtained by averaging over 10 random cross-validation data partitions. The vertical broken line represents mean CV-$R^2$ of the original data, which was 0.3901 (p-value = 0.0012).

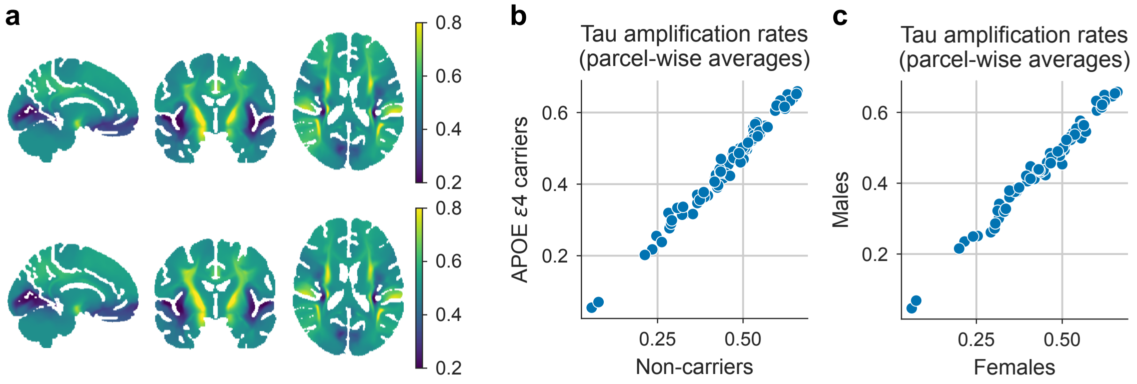


Supplementary Fig. 5: Stratification of inferred tau amplification rates by APOE genotype and sex. (a) Population-averaged maps of the inferred tau amplification rates. Top: APOE $\epsilon 4$ carriers; bottom: non-carriers. Sagittal, coronal, and axial slices are shown. (b) Parcel-wise comparison of amplification rates between APOE $\varepsilon 4$ carriers and non-carriers (Pearson $r = 0.991$). Brain parcellation was defined using the Desikan–Killiany atlas for cortical regions and FreeSurfer subcortical segmentation (aseg) for subcortical regions. (c) Parcel-wise comparison between males and females (Pearson $r = 0.992$). The same parcellation scheme was used as in (b).
